# WNT stimulation induced conformational dynamics in the Frizzled-Dishevelled interaction

**DOI:** 10.1101/2022.07.19.500578

**Authors:** Carl-Fredrik Bowin, Pawel Kozielewicz, Lukas Grätz, Maria Kowalski-Jahn, Hannes Schihada, Gunnar Schulte

## Abstract

Frizzleds (FZD_1-10_) are G protein-coupled receptors containing an extracellular cysteine-rich-domain (CRD) that presents the orthosteric binding site of the 19 mammalian WNTs, the endogenous agonists of FZDs. FZDs signal via a diverse set of effector proteins, of which Dishevelled (DVL1-3) is the most well studied and which acts as a hub for several FZD-mediated signaling pathways. However, the mechanistic details of how FZD-DVL interaction mediate pathway initiation and provide pathway selectivity remain an enigma. Here, we use bioluminescence resonance energy transfer-based assays employing Venus-tagged DVL2 together with NanoLuciferase-tagged FZD_5_ to investigate the WNT-3A- and WNT-5A-induced dynamics of the FZD-DVL interaction. Our biophysical assessment suggests that the ligand-induced BRET changes over time originate from conformational dynamics in the FZD_5_-DVL2 complex rather than recruitment dynamics of DVL2 to FZD_5_. Thus, we suggest that extracellular agonist and intracellular transducers could cooperate with each other through allosteric interaction with FZDs in a ternary complex reminiscent of that of classical GPCRs.

**One Sentence Summary:** Analysis of the interaction of FZD_5_ and DVL2 uncovers WNT-induced conformational dynamics of a WNT-FZD_5_-DVL2 complex.

## INTRODUCTION

The class F of G protein-coupled receptors (GPCRs) consists of ten Frizzled paralogues (FZD_1-10_) and Smoothened (SMO) (*1*). They bear the structural hallmarks of GPCRs, having an extracellular N-terminus, seven transmembrane spanning helices (TM1-7), three extracellular loops, three intracellular loops as well as an intracellular helix 8 and a C-terminus. FZDs also contain a highly conserved cysteine-rich domain (CRD) at the N-terminus constituting the binding site for their endogenous ligands, the lipoglycoproteins of the Wingless/Int1 (WNT) family (*2*). FZDs are subdivided into four homology clusters, FZD_1,2,7_, FZD_3,6_, FZD_4,9,10_ and FZD_5,8_ (*1*), which interact with 19 mammalian WNTs; however, family-wide ligand-receptor selectivity as well as ligand-receptor pathway selectivity remain poorly understood, although some light was recently shed on the complexity of WNT-FZD interaction (*3*).

FZDs recruit a diverse set of effector proteins (*4*). The phospho- and scaffold protein Dishevelled (DVL) is the most studied and acts as a hub for multiple intracellular WNT signaling pathways (*5–7*). Mammals express three DVL paralogues, DVL1-3, which consist of three distinct, structured domains: the N-terminal DIX (Dishevelled and Axin) domain, the central PDZ (Post-synaptic density protein-95, Disc large tumor suppressor, Zonula occludens-1) domain, the DEP (Dishevelled, Egl-10 and Pleckstrin) domain as well as a flexible C-terminus. The DIX domain, also found in the protein Axin, readily oligomerizes leading to the formation of dynamic, cytosolic DVL puncta (*8, 9*). While the DIX domain is essential for WNT/β-catenin signaling, where it forms dynamic DIX-DIX oligomers with other DVL or Axin proteins, it remains unclear how the DIX-DIX interaction integrates into the physical process of signal transduction initiated by WNT-activated FZD. The PDZ domain was initially found to interact with the conserved, but unconventional PDZ ligand KTXXXW localized in the helix 8 of FZDs (*10*). However, more recently the DEP domain has emerged to be the most important for FZD-DVL interaction (*11, 12*) and WNT/β-catenin signaling (*13*). The DEP domain is a compact ∼10.5 kDa domain and consists of three α-helices, a β-hairpin and two β-sheets, including a finger loop with K446 (aa numbering from human DVL2) located at the tip. Mutations in the DEP domain including the L445E and K446M mutation prevent FZD-DVL interaction, underlining the importance of this region for DVL recruitment to the plasma membrane and signal initiation (*12, 13*). In addition, DVL has multiple phosphorylation sites and becomes heavily phosphorylated in response to WNT signaling, detectable as an electrophoretic mobility shift (*14*). Kinases responsible for DVL phosphorylation include casein kinase 1δ and ε (CK1δ/ε), casein kinase 2 and protein kinase C (*5, 9, 15*).

While the concept of dynamic FZD-DVL interaction being relevant for WNT-induced and FZD-mediated signal transduction was formulated in the late 1990s (*16*), the recruitment dynamics of FZD-DVL interaction were notoriously difficult to investigate (i) because DVL forms dynamic puncta due to DIX-DIX oligomerization (*8*), (ii) overexpressed DVL strongly interacts with overexpressed FZD already in the absence of agonist stimulation (*17*) and (iii) cell lysis disrupts the FZD-DVL interaction (*18*). Thus, ligand-induced dynamics in FZD and DVL interaction could not be assessed systematically so far. Primarily, the extent of plasma membrane recruitment of DVL, as detected by confocal microscopy, has been used as a semi-quantitative assessment of its functionality and as an indirect measure of FZD interaction (*16*). In recent years, employment of total internal reflection fluorescence (TIRF) microscopy revealed that DVL2 investigated at endogenous levels is recruited to and oligomerizes at the plasma membrane in response to WNT-3A stimulation (*19*). Additionally, FZD_4_-DVL2 and FZD_6_-DVL2 interactions were investigated with bioluminescence resonance energy transfer (BRET) detecting increased BRET signal after stimulation with Norrin and WNT-5A, respectively (*20, 21*). Although these findings add important knowledge to the field, they do not provide mechanistic insights for FZD-DVL interaction.

The question whether the FZD-DVL interaction is dynamic presents a major gap in the current understanding of how DVL functions to transduce WNT-induced signaling downstream of FZDs. Therefore, we investigated the kinetics and dynamics of WNT-induced FZD_5_-DVL2 complex rearrangements by employing BRET sensors. This biophysical analysis in living cells revealed (i) FZD_5_-mediated increase in BRET signal between FZD and DVL2 (or DVL2-DEP) in response to WNT-3A and −5A, (ii) the importance of the DEP domain of DVL2 for FZD_5_-DVL2 recruitment and dynamics after WNT stimulation and (iii) an oligomerization-independent conformational change in the WNT-stimulated FZD_5_-DVL2 complex. Our data suggest that extracellular agonist and intracellular transducer cooperate with each other through an allosteric interaction with FZDs in a ternary complex reminiscent of that of classical GPCRs (*22*), where transmembrane allosteric cooperativity is essential for the interpretation of WNT binding towards DVL dynamics.

## RESULTS

### Venus-DVL2 is recruited to FZD_5_-Nluc

While we have used a bystander BRET setup with NanoLuciferase-tagged DVL2 (Nluc-DVL2) and a plasma membrane-anchored yellow fluorescent protein (Venus-KRas) together with FZDs to investigate ligand-independent, basal plasma membrane recruitment of DVL2 (*23*), this system was less suited to investigate the dynamic relationship between FZD and DVL upon WNT stimulation in more detail. This setup measures predominantly plasma membrane association. Therefore, we optimized a direct BRET setup based on an N-terminally tagged Venus-DVL2 and C-terminally tagged HA-FZD_5_-Nluc (FZD_5_-Nluc) (Fig. 1A), which could allow to distinguish recruitment from complex conformational changes. We first validated these fusion proteins by quantifying WNT-independent, basal recruitment of Venus-DVL2 to co-expressed FZD-Nluc by an acceptor titration approach (see Fig. S1A for surface expression of FZD_5_-Nluc). Therefore, a fixed amount of FZD_5_-Nluc was co-transfected with increasing amounts of Venus-DVL2 into HEK293T FZD_1-10_ knockout (ΔFZD_1-10_) cells. The BRET signal for FZD_5_-Nluc saturated with increasing expression of the acceptor construct (Fig. 1B). In this assay, we employed the β_2_ adrenoceptor (β_2_AR)-Nluc as a negative control. Here, increasing expression of the DVL2 acceptor construct resulted in a linear, non-saturable increase in the BRET signal indicative of random and unspecific interaction between these two proteins (*24, 25*). The titration curve allowed us to determine a suitable ratio of expression levels between Venus- and Nluc-tagged constructs for further experiments, where saturation of DVL2 recruitment to FZD_5_ in the basal state is important to ensure consistency between experiments. Additionally, the basal recruitment of DVL by FZDs is independent of endogenously produced WNTs (*17, 21*). In order to validate the constitutive, ligand-independent DVL2 recruitment by FZD_5_ co-expression also in the current set-up we used the porcupine inhibitor C59 (Fig. S1B). Porcupine inhibitors prevent the maturation and secretion of WNT proteins and allow thereby to create conditions with severely reduced autocrine WNT input (*26*). Since no differences in BRET between FZD_5_ and DVL2 were detected in the absence or presence of C59, we did not include the porcupine inhibitor in subsequent experiments.

**Fig. 1.**
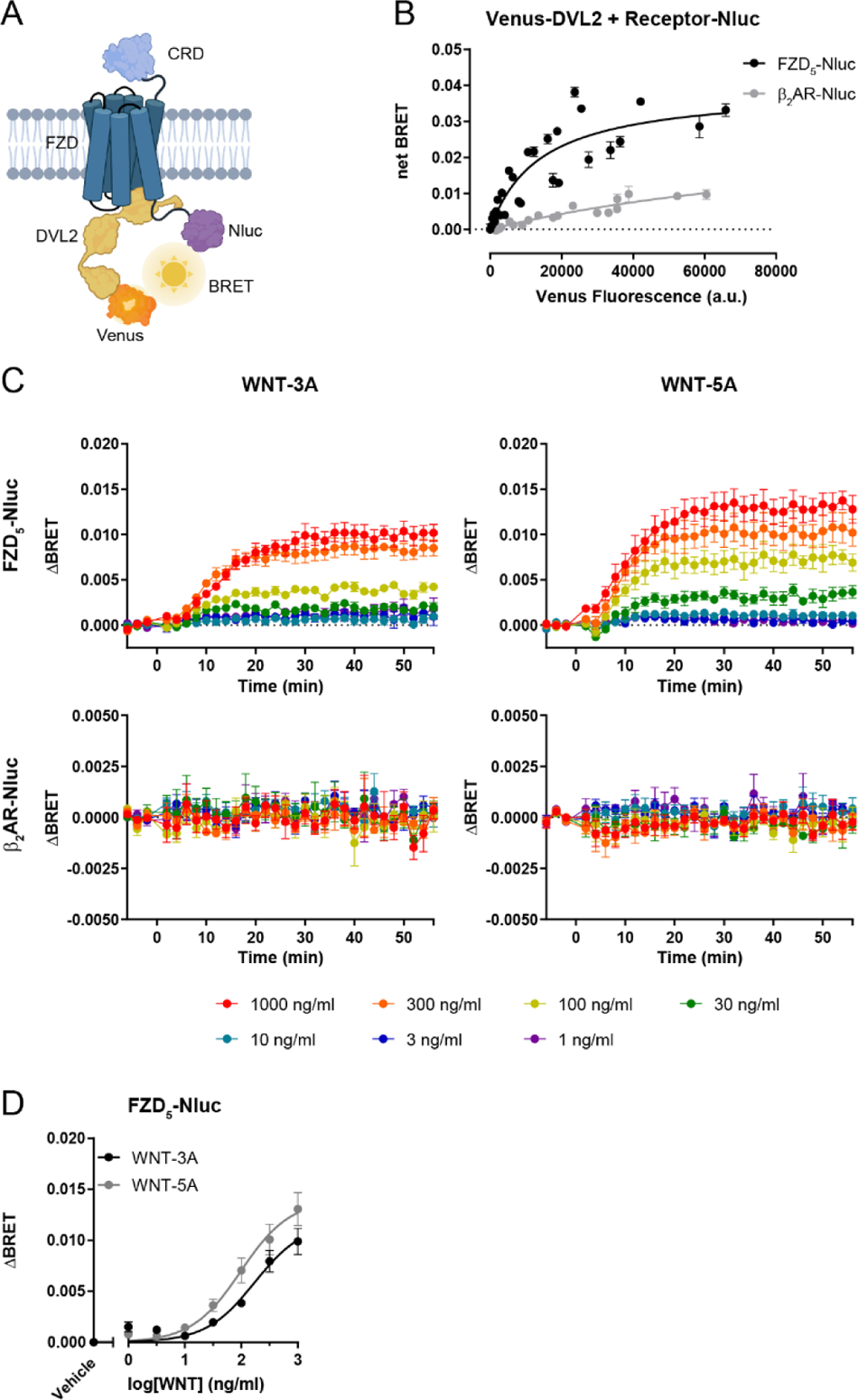
Venus-DVL2 is recruited to FZD_5_-Nluc and WNT stimulation induces FZD-DVL dynamics. (**A**) Schematic of the Venus-DVL2 and FZD_5_-Nluc BRET assay setup. DVL2 in yellow, Venus fused to the DVL2 N-terminal in orange and Nluc fused to the FZD_5_ C-terminus in purple. The scheme was prepared with the web-based tool BioRender.com. (**B**) Venus-DVL2 was titrated with a fixed amount of Nluc-tagged FZD_5_ in ΔFZD_1-10_ cells to assess WNT-independent, basal recruitment of DVL2 to FZD_5_; β_2_AR was used as negative control. net BRET is presented as mean ± SD of 3-5 independent experiments. (**C**) Fixed amounts of Venus-DVL2 and Nluc-tagged FZD_5_ (Venus:Nluc ratio 25:1) were transfected into ΔFZD_1-10_ cells to assess the change in FZD-DVL dynamics upon WNT stimulation. The kinetic BRET response between Venus-DVL2 and Nluc-FZD_5_ was monitored with WNT-3A or WNT-5A stimulation of FZD_5_ or the negative control β_2_AR. (**D**) Concentration response curve for WNT stimulation of FZD_5_-transfected cells based on BRET values 30 min after stimulation. ΔBRET values are presented as mean ± SEM of 3-7 independent experiments.

### WNTs induce a concentration-dependent change in FZD_5_-DVL2 interaction

With the aim to quantify WNT-induced dynamics of FZD_5_-DVL2 interaction, we monitored BRET between Venus-DVL2 and Nluc-tagged FZD_5_ in transiently transfected ΔFZD_1-10_ cells. After generating a baseline of the BRET signal, the cells were stimulated with increasing concentrations of either recombinant WNT-3A or WNT-5A (Fig. 1C). WNT stimulation in FZD_5_-Nluc transfected cells resulted in a concentration-dependent increase in the BRET signal over time for both WNT-3A and WNT-5A reaching a plateau for the high and intermediate concentrations of WNT. Again, β_2_AR-Nluc served as negative control, showing neither an increase nor a decrease in the BRET response to WNT-3A or WNT-5A treatment.

The data were summarized to quantify the concentration dependency of the WNT-induced FZD_5_-DVL dynamics. For that purpose, we plotted the signal response 30 min after stimulation to acquire a concentration response curve (Fig. 1D), which resulted in a sigmoidal curve for both WNTs, with an EC_50_ value for WNT-3A of 230.9 ng/ml (6.2 nM) [95% CI 110.1-484.6 ng/ml (2.9-13.0 nM)] and for WNT-5A of 117.9 ng/ml (3.1 nM) [95% CI 58.6-237.3 ng/ml (1.5-6.2 nM)]. When comparing the WNT-3A- and the WNT-5A-elicited responses, no statistically significant difference in the maximum ΔBRET response was observed.

To control that the WNT-induced BRET response was indeed elicited by the added WNT proteins, we heat-inactivated WNT-3A. The inactivated WNT preparation did not induce an increase in the BRET response observed between Venus-DVL2 and FZD_5_-Nluc (Fig. S1C). We also addressed the potential importance of endogenously expressed DVL, which could compete with exogenous Venus-DVL2 and thereby affect the BRET response. In order to investigate the role of endogenously expressed DVL1, 2, 3, we used triple knock out HEK293 cells devoid of DVL1, 2, 3 (ΔDVL1-3) (*13*). However, in the absence of endogenous DVL, the WNT-induced BRET signal between FZD_5_-Nluc and Venus-DVL2 still resembled the kinetics observed in ΔFZD_1-10_ cells (Fig. S1D). While FZD_5_ expression results in a ligand-independent, constitutive recruitment of DVL2, the question remains, whether the observed WNT-induced ΔBRET reports additional recruitment of DVL molecules to the surface-expressed FZDs (recruitment dynamics) or rather a conformational rearrangement in the preformed FZD_5_-DVL2 complex (conformational dynamics). In order to exclude the former, we used a bystander BRET set-up between the Nluc-DVL2 and membrane-anchored Venus (Venus-KRas) in the presence of untagged FZD_5_, similar to what we have published previously (Fig. 2A) (*23, 27*). If WNT stimulation of FZD_5_ would lead to additional recruitment of Nluc-DVL2 to the membrane, the bystander BRET would report an increased proximity of Nluc-DVL2 to Venus-KRas. However, neither WNT-3A nor WNT-5A stimulation (1 µg/ml), which resulted in a substantial ΔBRET in the direct BRET approach shown in Fig. 1C, elicited any changes in the bystander BRET read-out (Fig. 2B). Thus, these data support the hypothesis that the observed BRET changes in the direct BRET set-up originate from complex rearrangement (conformational dynamics) rather than additional recruitment of new DVL molecules to membrane-embedded FZDs (recruitment dynamics). To validate the ability of the bystander BRET assay to measure changes in DVL2 membrane recruitment, we used digitonin-induced membrane permeabilization. While basal bystander BRET recordings prior digitonin reported on the constitutive FZD_5_-mediated DVL2 recruitment, the decrease upon digitonin treatment and the lower BRET values subsequent to permeabilization emphasize the sensitivity of the approach to record plasma membrane recruitment of DVL2 (Fig. 2C).

**Fig. 2.**
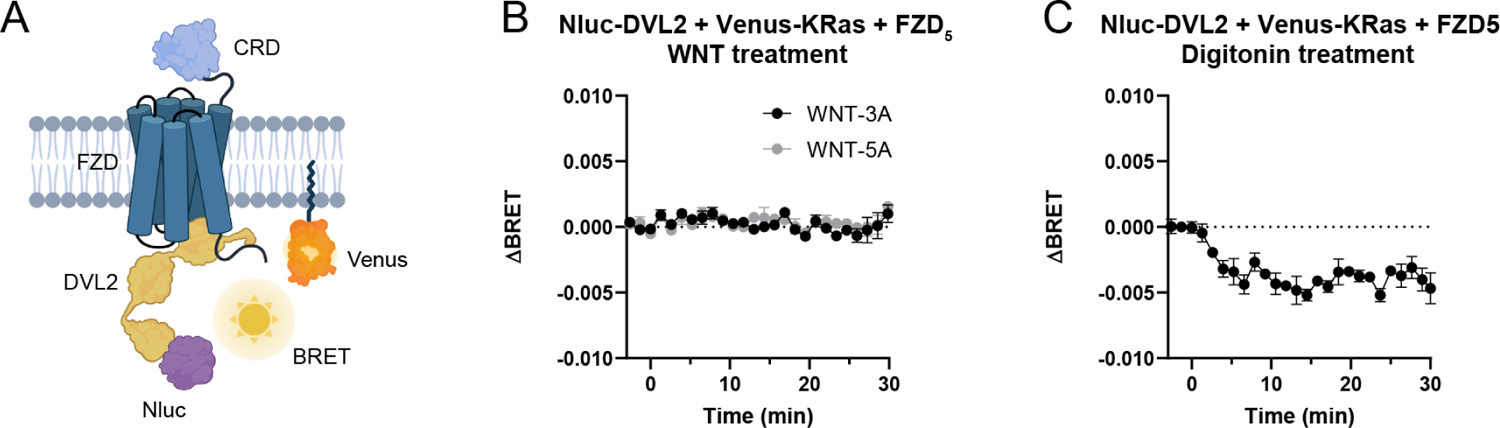
Bystander BRET monitoring FZD_5_-induced DVL recruitment to the membrane supports WNT-induced conformational over recruitment dynamics. (**A**) Schematic of the Nluc-DVL2 and Venus-KRas bystander BRET assay. Membrane-anchored Venus-KRas in yellow, Nluc fused to the DVL2 N-terminal in orange. The scheme was prepared with the web-based tool BioRender.com. The plasma membrane recruitment bystander BRET between Nluc-DVL2 and Venus-KRas was measured in response to either (**B**) 1 µg/ml of WNT-3A or WNT-5A and (**C**) 10 µg/ml of digitonin. Data show mean ± SEM of three independent experiments.

### Oligomerization-deficient DVL2 responds to WNT stimulation

While full activation of the WNT/β-catenin signaling pathway requires the functional DIX domain of DVL to allow for the accumulation of β-catenin (*28*), we hypothesized that the WNT-induced dynamics in the FZD-DVL interaction assessed by BRET is independent of DVL2 oligomerization. The mutant DVL2-M2/M4 harbors two mutations in the DIX domain resulting in an oligomerization-deficient protein (*8*). Thus, we used a Venus-DVL2-M2/M4 construct to assess the role of DVL-DVL oligomerization for the FZD-DVL interplay. First, we validated basal interaction of Venus-DVL2-M2/M4 with FZD_5_-Nluc in a BRET acceptor titration experiment. Despite the lower expression levels of Venus-DVL2-M2/M4 compared to wildtype Venus-DVL2 we observed an expression-dependent, saturable and thus specific interaction with FZD_5_ but not β_2_AR (Fig. 3A). We then investigated the WNT-induced FZD-DVL dynamics with this mutant as done previously with the wildtype Venus-DVL2. The WNT-3A- and WNT-5A-induced kinetic responses for Venus-DVL2-M2/M4 and FZD_5_-Nluc were similar to the results obtained with wildtype Venus-DVL2 (Fig. 3B). In order to determine the WNT-induced rate constant k of the conformational dynamics within the FZD-DEP complex, we fitted each individual experiment with a plateau followed by one phase association equation (Fig. S2A-B). No significant differences were found when comparing the respective WNT-induced conformational dynamic rate constant, k, between Venus-DVL2 and Venus-DVL2-M2/M4 (Fig. 3C), emphasizing that oligomerization hardly contributes to the observed conformational dynamics.

**Fig. 3.**
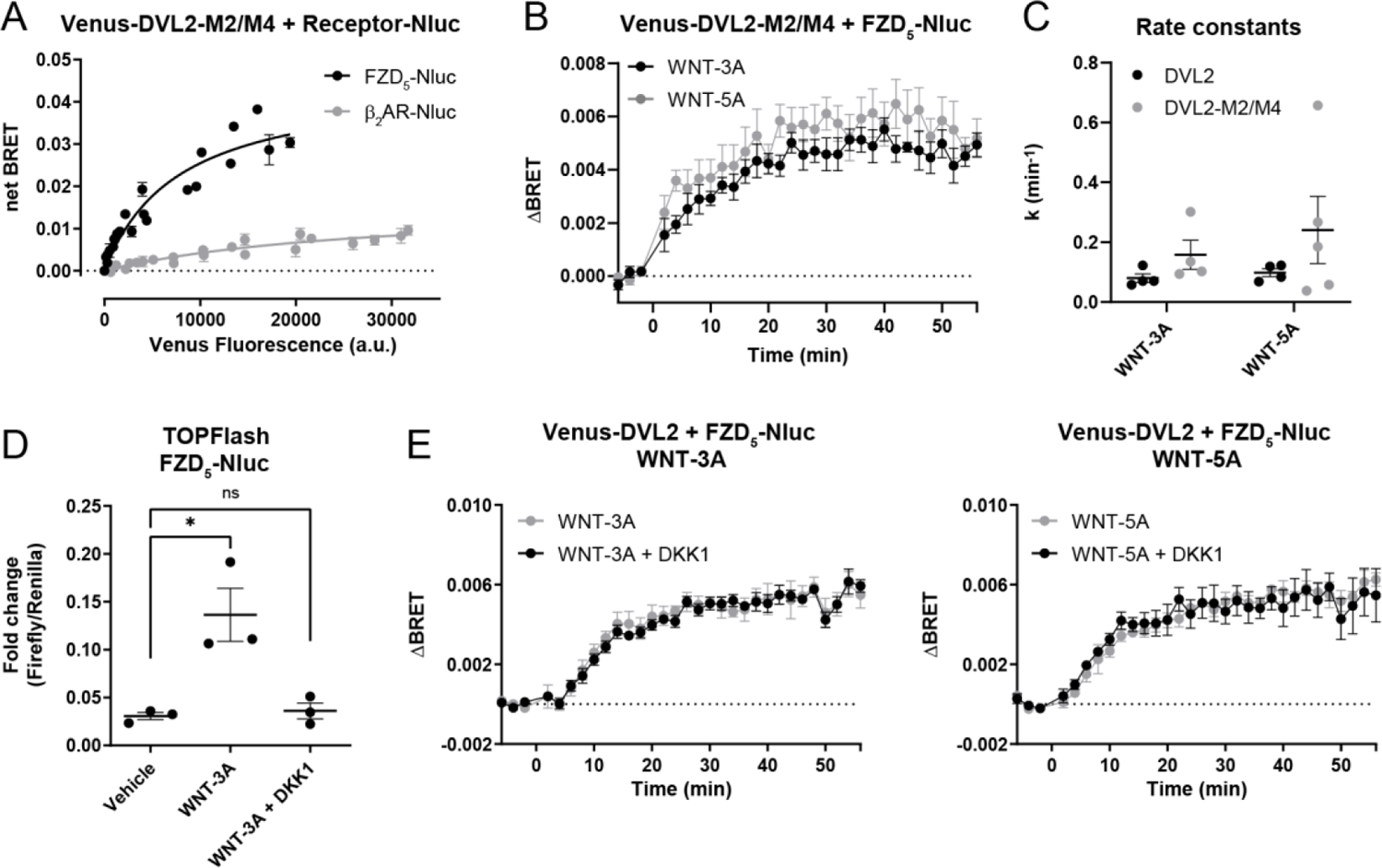
FZD-DVL dynamics can be separated from DVL-oligomerization and signalosome formation. (**A**) Venus-DVL2-M2/M4 was titrated with a fixed amount of FZD_5_-Nluc in ΔFZD_1-10_ cells to assess basal recruitment of oligomerization-impaired DVL2; β_2_AR was used as negative control. (**B**) The kinetic BRET response between Venus-DVL2-M2/M4 and FZD_5_-Nluc (Venus:Nluc ratio 25:1) upon 1 µg/ml of WNT-3A or WNT-5A stimulation was monitored. Data show mean ± SEM of 4-5 independent experiments. (**C**) Comparison of Venus-DVL2 and Venus-DVL2-M2/M4 rate constants k upon WNT stimulation. No statistically significant differences were found between DVL2 and DVL2-M2/M4 (two-way ANOVA with Fisher’s LSD post hoc analysis). Data presented in (C) are extracted from the curve fitting presented in Fig. S2A-B. (**D**) The bar graph depicts the TOPFlash response in ΔFZD_1-10_ cells transfected with FZD_5_-Nluc after 24 h of stimulation with either vehicle or 300 ng/ml of WNT-3A in the absence of presence of 300 ng/ml of DKK1. Data are presented as mean ± SEM of 3 independent experiments (* P < 0.05, ns = not significant, one-way ANOVA with Dunnett’s post hoc analysis). (**E**) The kinetic BRET response between Venus-DVL2 and FZD_5_-Nluc (Venus:Nluc ratio 25:1) was monitored in ΔFZD_1-10_ cells during stimulation with 300 ng/ml of either WNT-3A or WNT-5A in the absence or presence of 300 ng/ml of DKK1. ΔBRET is presented as mean ± SEM of 4 independent experiments.

### FZD-DVL dynamic response is independent of LRP5/6

DVL is a central mediator of β-catenin-dependent and -independent WNT signaling; however mechanistic details remain unclear (*5*). One of the current working models for signal perception and pathway initiation is based on a signalosome mechanism, where WNTs bind simultaneously to both FZDs and low-density lipoprotein receptor-related protein 5/6 (LRP5/6) initiating DVL/Axin oligomerization (*29–31*). While this concept appears valid for WNTs that activate the WNT/β-catenin pathway, WNT-5A, which generally activates β-catenin-independent signaling, also binds LRP5/6 without feeding into the WNT/β-catenin pathway (*32*). In order to address the role of LRP5/6 as WNT co-receptors and their involvement in mediating the agonist-induced FZD-DVL dynamics, we employed recombinant Dickkopf-related protein 1 (DKK1), which acts as a WNT/β-catenin signaling pathway inhibitor by binding LRP5/6, preventing the formation of the signalosome complex (*33, 34*). Therefore, we employed our BRET setup using Venus-DVL2 and FZD_5_-Nluc in ΔFZD_1-10_ cells and treated them with 300 ng/ml of either WNT-3A or WNT-5A in the absence or presence of 300 ng/ml of recombinant DKK1. While the DKK1 concentration used in these experiments abrogated WNT-3A-induced WNT/β-catenin signaling (Fig. 3D), the addition of DKK1 did not affect the BRET response induced by WNT-3A or WNT-5A (Fig. 3E), clearly separating LRP5/6-dependent signalosome formation from the WNT-induced FZD_5_-DVL2 dynamics observed in the BRET assay.

To further manifest the independency of the WNT-induced FZD-DVL dynamics from LRP5/6, we used HEK293 cells devoid of LRP5/6 (ΔLRP5/6). In order to assess ligand-induced effects, we transiently transfected ΔLRP5/6 cells with Venus-DVL2 and FZD_5_-Nluc and monitored the BRET signal before and after stimulation with WNT-3A or WNT-5A. These experiments revealed a similar response pattern with saturating curves and similar rate constants, despite an overall weaker ΔBRET signal (Fig. S3A-C). The weaker response in the ΔLRP5/6 cells likely originated from the lower transfection efficiency, which is visualized as values for luciferase and Venus counts in comparison to ΔFZD_1-10_ cells (Fig. S3D). Taken together, these data argue that WNT-induced FZD_5_-DVL2 dynamics are independent of LRP5/6 and could precede formation of the WNT-3A-induced signalosome.

### The DEP domain of DVL responds to WNT stimulation

During recent years, the DEP domain has emerged as the primary interaction site between DVL and FZDs (*12, 13*). This statement is further corroborated by an acceptor titration BRET experiment employing the Venus-tagged DVL2 lacking the DEP domain (ΔDEP DVL2) in combination with Nluc-tagged FZD_5_ or β_2_AR (negative control). While FZD_5_ recruited full length DVL2 (Fig. 1B), the BRET amplitude in the presence of FZD_5_-Nluc with increasing levels of ΔDEP Venus-DVL2 was not different from that of the negative control (Fig. S4A), asserting the DEP domain as the major FZD-interaction site (*16*). As previously surmised (*4*), we wanted to employ a minimal DVL domain to address the question whether the complexity of FZD-DVL interaction and most importantly their ligand-induced conformational dynamics can be reduced to the minimal DEP domain, serving as a conformational sensor for basal and WNT-activated FZDs. For this purpose, we fused Venus to the C-terminus of the DEP domain (aa S418-G510) of human DVL2 (DEP-Venus) (Fig. 4A; see also Supplementary Materials & Methods). Titration of DEP-Venus with a fixed amount of FZD_5_-Nluc (Fig. 4B) emphasized saturable and specific interaction with FZD_5_-Nluc already at very low expression levels, indicating high affinity between FZD_5_ and DEP, in line with previous findings employing microscopy-based assessment of FZD-DEP interaction (*11, 12, 17*). Furthermore, both WNT-3A and WNT-5A elicited a dynamic BRET response between FZD_5_-Nluc and DEP-Venus (Fig. 4C) and the response for FZD_5_-Nluc reached a plateau after stimulation with both WNT-3A and WNT-5A. The concentration response for FZD_5_-Nluc also reflected what we observed with Venus-DVL2, with an EC_50_ value for WNT-3A of 110.1 ng/ml (2.9 nM) [95% CI 49.9-243.1 ng/ml (1.3-6.5 nM)] and for WNT-5A of 277.5 ng/ml (7.3 nM) [95% CI 150.7-511.1 ng/ml (4.0-13.5 nM)] (Fig. 4D). Intriguingly, there was a more pronounced difference between the WNT-3A and WNT-5A response, indicative of distinct WNT subtype-dependent complex formation between FZD_5_ and DEP. Additionally, we used inactivated WNT-3A, which was unable to elicit a BRET response, as a control (Fig. S4B). Also, there was no difference in the rate constants of the WNT-induced kinetic ΔBRET response of DVL2 and DEP (Fig. 4E, S5A), further asserting DEP as a conformational sensor of FZD-DVL interaction and dynamics.

**Fig. 4.**
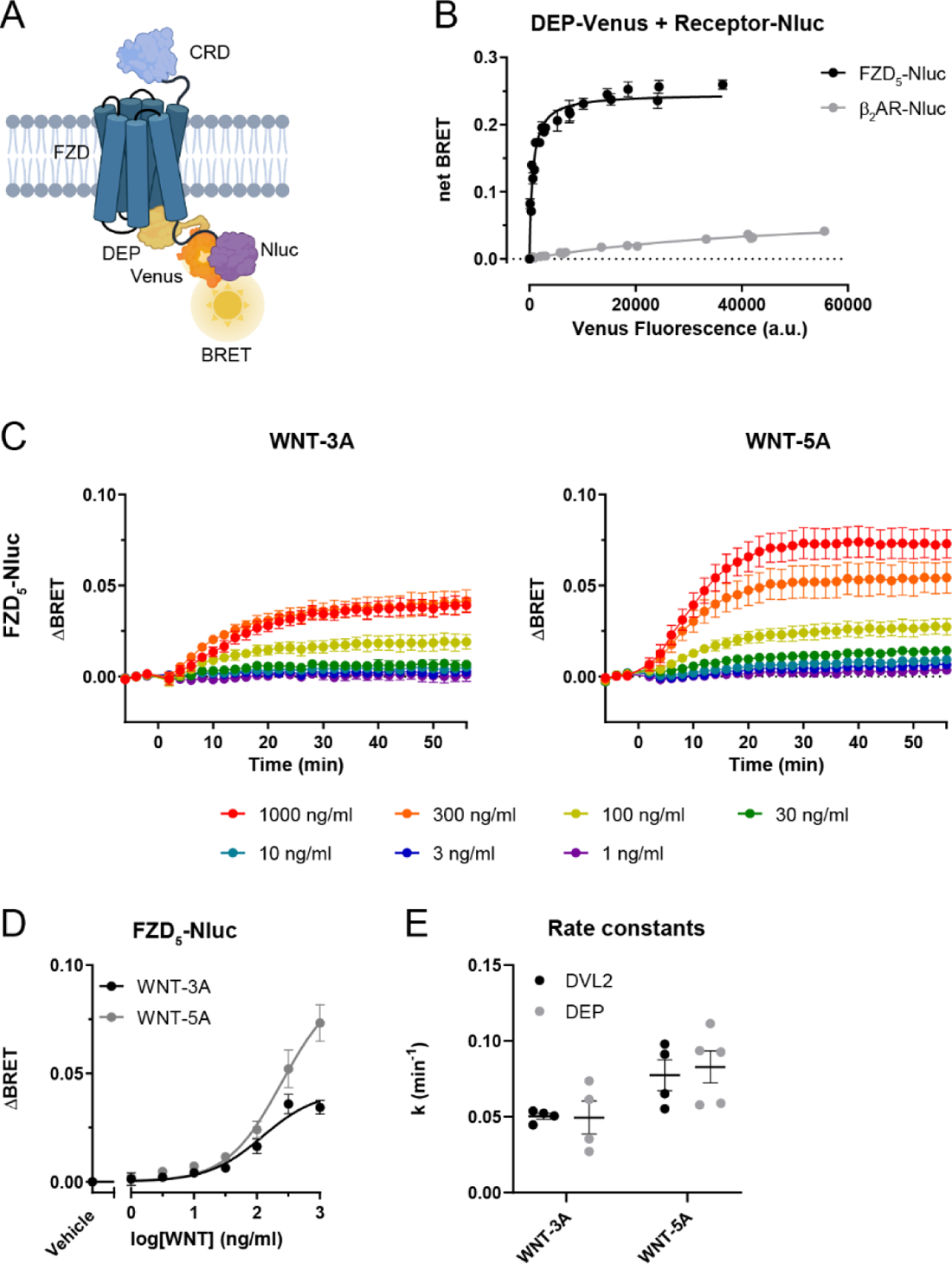
DEP-Venus is a conformational sensor for an active FZD_5_. (**A**) Schematic presentation of the DEP-Venus and FZD_5_-Nluc BRET assay setup. DEP in yellow, Venus fused to the DEP C-terminus in orange and Nluc fused to the FZD_5_ C-terminus in purple. The scheme was prepared with the web-based tool BioRender.com. (**B**) DEP-Venus was titrated with a fixed amount of FZD_5_-Nluc in ΔFZD_1-10_ cells to assess basal recruitment to FZD_5_; β_2_AR was used as a negative control. (**C**) The kinetic BRET response was monitored between DEP-Venus and FZD_5_-Nluc (Venus:Nluc ratio 25:1) in ΔFZD_1-10_ cells during WNT-3A or WNT-5A stimulation. Data show mean ± SEM of 3 independent experiments. (**D**) Concentration response curve for FZD_5_-mediated effects is based on data obtained 30 min after stimulation. ΔBRET is presented as mean ± SEM of 3 independent experiments. (**E**) Comparison of Venus-DVL2 and DEP-Venus rate constants k upon WNT stimulation. No statistical differences were found between DVL2 and DEP (two-way ANOVA with Fisher’s LSD post hoc analysis). Data presented in (E) are extracted from the curve fitting presented in Fig. S2A and Fig. S5A.

### Mutational analysis of FZD-DEP dynamics

To investigate the FZD-DEP interaction and BRET conformational dynamics in more detail, we employed previously described DEP mutants (Fig. 5A) (*11–13*). While these mutants have previously been investigated for their recruitment to FZD_5_ by confocal microscopy in the absence of WNT stimulation, we aimed here to employ kinetic BRET experiments to better understand the impact of DEP mutations on WNT-induced conformational dynamics in a FZD_5_-DEP complex. The G436P mutant was reported to be impaired in its ability to form DEP dimers, allowing us to address the role of DEP dimers for the WNT-induced FZD-DEP dynamics. The K446M mutant, localized at the tip of the DEP finger loop, was reported to block recruitment of DEP and full length DVL2 to FZD_5_, potentially pinpointing an interaction site between FZD and DEP. Lastly, the L445E mutant – adjacent to K446M in the DEP finger loop – was also reported to be unable to bind FZD_5_ both as a DEP construct and full length DVL2. All these mutations in full length DVL2 impair WNT/β-catenin signaling (*13*). First, we assessed the basal recruitment of the three mutants to FZD_5_- and β_2_AR-Nluc (the latter used as a negative control) with BRET acceptor titrations (Fig. 5B-D). While we could confirm that the G436P DEP dimer mutant interacts with FZD_5_, albeit with a lower saturation BRET_max_ and reduced BRET_50_ as compared to wildtype DEP (Fig. 5B, E), we could also detect recruitment of the K446M mutant to FZD_5_. This interaction, however, also presented with a reduced saturation BRET_max_ and BRET_50_ (Fig. 5C, E). Furthermore, we did not detect any specific interaction between FZD_5_ and the DEP L445E mutant, corroborating previous results (Fig. 5D, E) (*11*). Therefore, we continued to investigate the WNT-induced FZD_5_-DEP conformational dynamics with the FZD-interacting G436P and K446M DEP mutants (Fig. 6A-B). They both presented with conformational dynamic FZD-DEP ΔBRET responses upon WNT-3A and WNT-5A stimulation. Thus, the dimer deficient DEP G436P mutant and the finger loop K446M mutant maintain the conformational dynamic BRET response over time, albeit with a different kinetic profile. In order to determine kinetic parameters such as a the WNT-induced rate constant k and BRET_max_ of the conformational dynamics within the FZD-DEP complex, we fitted each individual experiment with a plateau followed by one phase association equation (Fig. S5B-C). It should be noted that the comparison of kinetic BRET_max_ between mutants is compromised by their differences in the saturation BRET_max_ values reported in Fig. 5E. On the other hand, rate constants can be compared between the groups. Thus, the assessment of FZD-DEP interaction by BRET extends our possibilities to distinguish constitutive (ligand-independent) recruitment of DEP to FZD and ligand-elicited conformational dynamics suggesting conformational rearrangement in the FZD-DEP interface as a hallmark of WNT/FZD signal initiation. We found that the DEP G436P mutant presented with a significantly increased kinetic rate with both WNT-3A and WNT-5A stimulation compared to wildtype DEP (Fig. 6C). Further, the DEP K446M mutant showed an increased kinetic rate with WNT-5A stimulation compared to wildtype DEP. Comparing WNT-3A and WNT-5A induced kinetic ΔBRET_max_, we observed a difference for both wildtype DEP and DEP G436P, but not for DEP K446M (Fig. 6D).

**Fig. 5.**
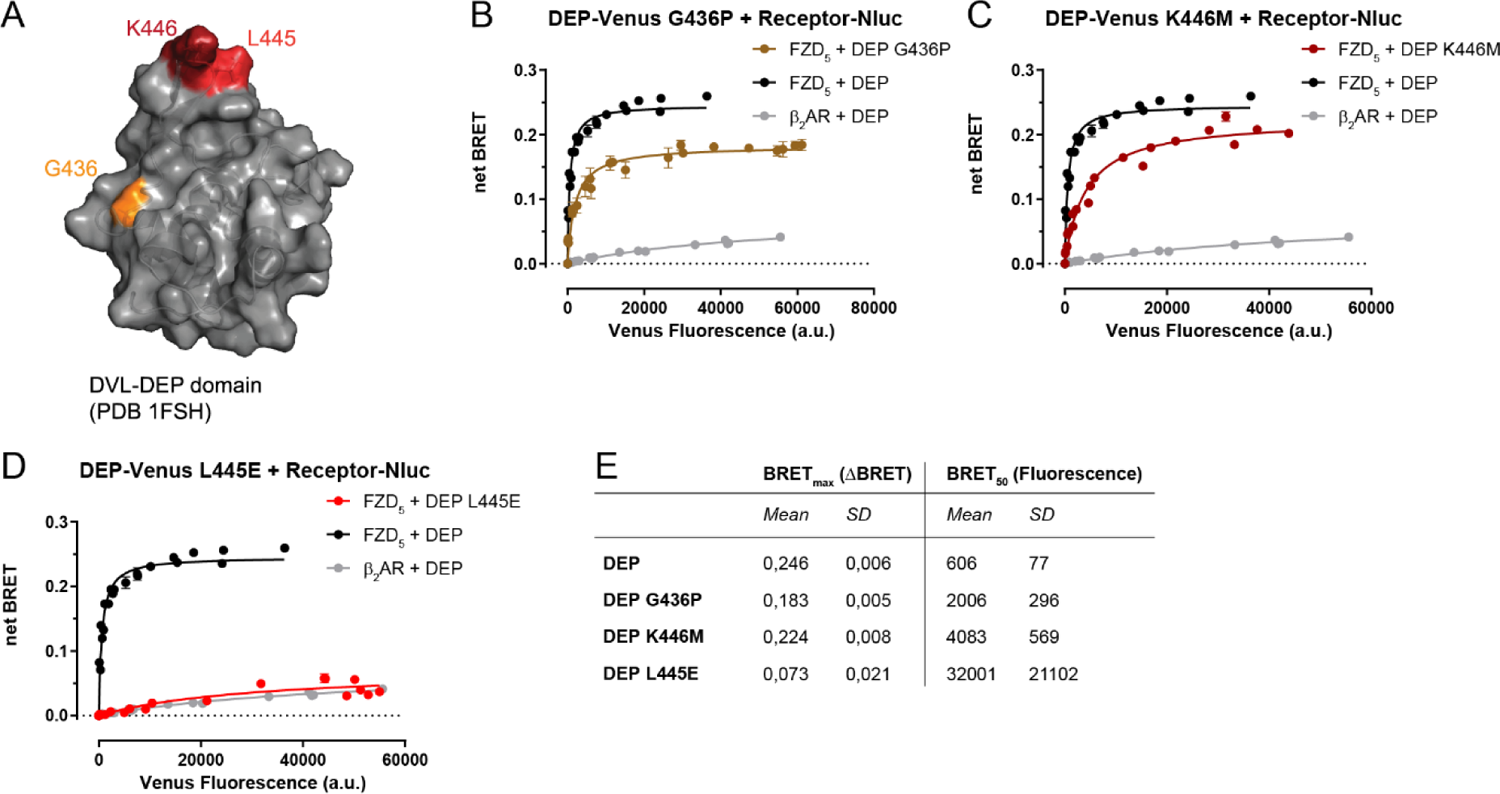
Mutations in the DEP domain change FZD_5_-DEP interaction. (**A**) The DEP mutations G436P, K446M and L445E (numbering for human DVL2) are visualized in a structure of the DEP domain of mouse DVL1 (PDB: 1FSH). (**B-D**) Titration of either DEP-Venus mutant (B) G436P, (C), K446M or (D) L445E with a fixed amount of FZD_5_-Nluc in ΔFZD_1-10_ cells to assess basal recruitment; β_2_AR was used as negative control (same as in Fig. 4B). Data show mean ± SD of three independent experiments. (**E**) Summary of saturation BRET_max_ and BRET_50_ values of wildtype DEP and mutants extracted from fits in Fig. 4B and Fig. 5B-D.

**Fig. 6.**
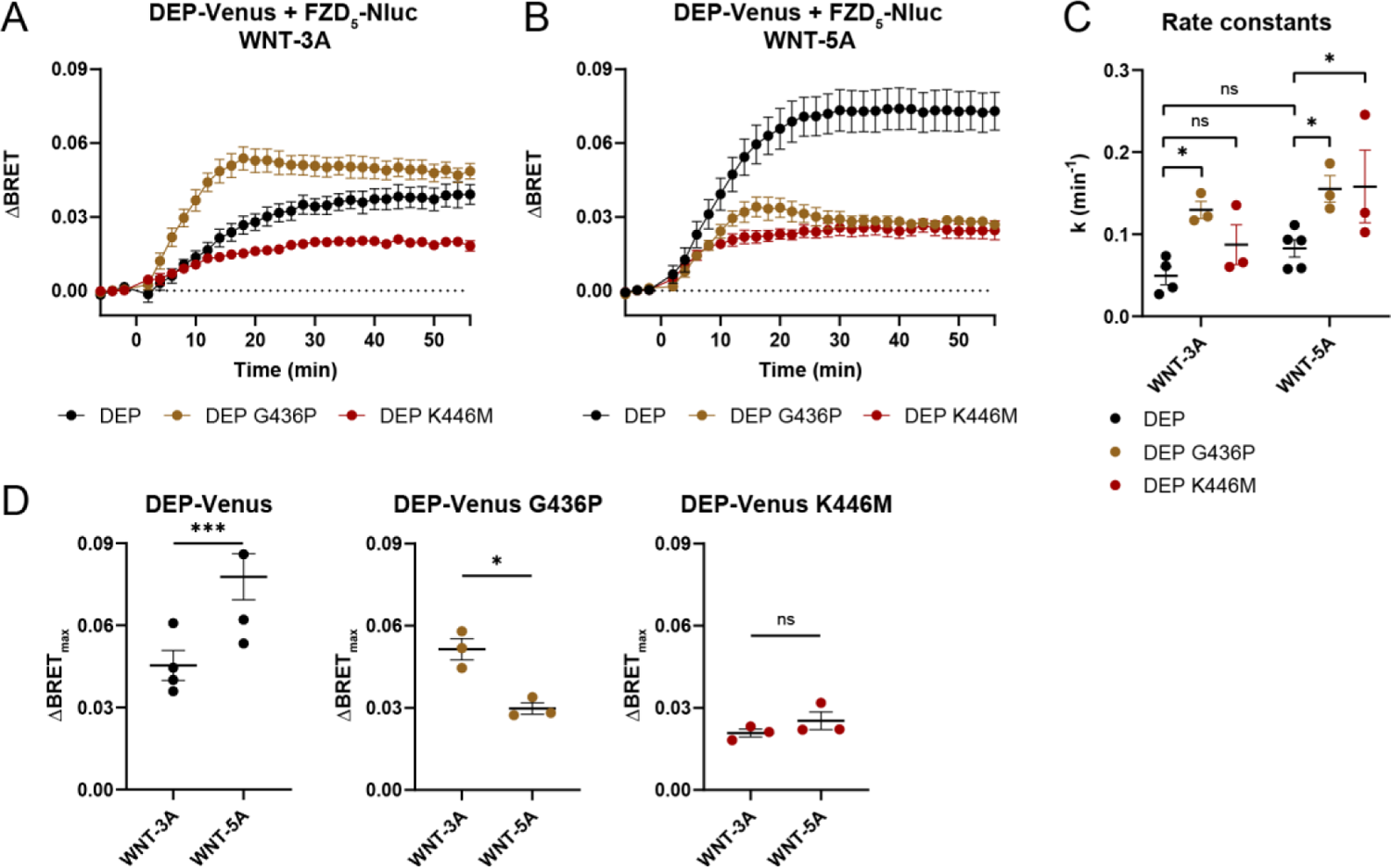
Mutations in the DVL DEP domain affect agonist-induced conformational dynamics in the FZD_5_-DEP interplay. (**A-B**) The kinetic BRET response was monitored between DEP-Venus G436P or DEP-Venus K446M and FZD_5_-Nluc in ΔFZD_1-10_ cells during (A) WNT-3A or (B) WNT-5A stimulation. Data show mean ± SEM of 3 independent experiments. DEP-Venus data is from Fig. 4B. (**C**) Comparison of DEP-Venus constructs rate constants k upon WNT stimulation (significance levels are given as * P < 0.05 or ns (not significant), two-way ANOVA with Fisher’s LSD post hoc analysis). (**D**) ΔBRET_max_ values for the different DEP constructs with WNT stimulation. Observe that comparisons of conformational BRET_max_ values between mutants are not quantitative provided the different saturation BRET_max_ values obtained in acceptor titration experiments shown in Fig. 5E. Data presented in (C) and (D) are extracted from the curve fitting presented in Fig. S5A-C.

## DISCUSSION

DVL plays a central role in both β-catenin-dependent and -independent WNT signaling but the mechanistic contribution of constitutive recruitment and ligand-induced dynamics in the FZD-DVL interplay remains obscure. While WNT-induced recruitment dynamics, referring to the recruitment of cytosolic DVL to FZD, have been postulated since the discovery of FZD-mediated DVL recruitment to the plasma membrane, a suitable quantitative assay methodology has so far not been available to investigate FZD-DVL dynamics. As pointed out already in 2012, understanding the dynamic interplay between FZD and DVL will provide essential mechanistic details to grasp FZD-mediated WNT pathway selectivity (*12*). Here, we fill this knowledge gap and employ a sensitive BRET approach to shed light onto the WNT-elicited conformational rather than recruitment dynamics in the FZD-DVL interaction in living cells. While this approach is suitable to study the interplay between FZD and full length DVL, we also explore the use of a minimal DEP domain as a BRET partner, which generally recapitulates the WNT-induced FZD-DVL conformational dynamics and allows to decipher important details in this interface at the crossroads of the WNT signaling system. It should also be noted that this experimental approach is suitable to assess WNT-induced dynamics in the interplay of DVL with other proteins, such as the transmembrane proteins ROR1/2, RNF43 or LRP5/6, to name but a few.

The DEP domain of DVL is important for signal transduction (*13*) and for FZD-DVL interaction in the basal state (*11, 12, 16, 17*). With the help of TIRF microscopy, it was previously shown that WNT-3A stimulation has an effect on DVL oligomerization and its recruitment to the plasma membrane (*19*). In addition, direct BRET was also employed to assess the effect of Norrin stimulation on FZD_4_-DVL2 interaction (*20*). Despite these recent advances, contradictory results point in opposite directions. While the above-mentioned results advocate an increase in FZD-DVL recruitment and DVL-DVL oligomerization at the plasma membrane, it was also concluded that the FZD_5_-DEP interaction decreases in response to WNT-3A stimulation (*11*) and that WNT-3A stimulation had no effect on DVL oligomerization or plasma membrane recruitment (*35*). Our BRET analysis allows to resolve molecular details behind FZD-DVL interaction dynamics, and we conclude that acute WNT stimulation affects the constitutively formed FZD-DVL complex. As BRET depends upon both the distance and dipole orientation between acceptor and donor (*36*), direct BRET data could be interpreted either as a change in FZD_5_-DVL2 recruitment or an alteration in the overall conformation of the FZD_5_-DVL2 complex. However, provided (i) that the experimental conditions favor FZD-DVL saturation prior to WNT stimulation, (ii) the absence of a response in the bystander BRET set up (Fig. 2C) and (iii) the DVL2-M2/M4 oligomerization-deficient mutant still induced a BRET response (Fig. 3B), we argue that the reported WNT-induced ΔBRET response reflects conformational dynamics in a pre-formed FZD-DVL or FZD-DEP complex. Given the similar BRET profiles and rate constants in response to WNTs using either wildtype DVL2 or DVL2-M2/M4, we can separate WNT-induced FZD-DVL dynamics from WNT-induced DVL-DVL oligomerization. Furthermore, because there is a substantial basal FZD-DVL interaction and the BRET assays are performed at the saturated end of the spectrum, meaning that the equilibrium is heavily shifted towards basal FZD-DVL interaction, it is reasonable to assume that a large part of the membranous FZD population is already bound to DVL2. As a consequence, in the present experimental system acute WNT stimulation does most likely not lead to a significant increase in DVL recruitment to yet unbound FZDs, supported by our bystander BRET data (Fig. 2C). Therefore, we interpret the WNT-induced ΔBRET as a rearrangement in the pre-existing FZD-DVL complex, which implicates a WNT-induced conformational change in the core of FZD, analogous to ligand-controlled, allosteric receptor/transducer coupling known from other GPCR families (*4*). Thus, it is of utmost importance to underline that the experimental conditions used in our work, where most surface expressed FZDs are bound to a DVL (or DEP) molecule, allow us to differentiate ligand-induced recruitment dynamics from conformational dynamics in the FZD/DVL or FZD/DEP complex. We surmise that DVL dynamics in cells with physiological DVL expression levels present as a composite response consisting of WNT-induced recruitment and conformational dynamics.

The observation that the minimal DEP domain recapitulates the WNT-induced dynamics observed with full length DVL2, including similar rate constants (Fig. 4E), argues that the DEP domain serves as a conformational sensor of a FZD conformation that is active with regard to DVL interaction. This is particularly relevant in the light of previous findings showing that the mutation of the conserved molecular switch in TM6 and TM7 of FZDs improves the likelihood of the agonist bound FZD_5_ to recruit a mini-G protein at the same time as it abrogates the basal recruitment of DVL2 to FZD_5_ (*23*). In combination, these findings argue for a conformational selection of transducer interaction, where distinct FZD conformations are required to feed into DVL vs G protein interaction and signaling. Moreover, the difference in the WNT-3A- and WNT-5A-induced FZD_5_-DEP BRET response is intriguing, where we observed that WNT-5A elicited a larger kinetic ΔBRET_max_ compared to WNT-3A (Fig. 6D). Two likely explanations for this are that (i) the two WNTs stabilize different receptor conformations or (ii) they stabilize the same conformation but with different probabilities (*37, 38*).

In order to better understand the general importance of FZD-DVL dynamics for WNT-induced FZD signaling, we have chosen FZD_5_, which is intensively studied and which both initiates the WNT/β-catenin pathway and interacts with and activates heterotrimeric G proteins in a WNT-dependent fashion (*12, 13, 23, 39, 40*).

In an effort to further dissect the observed WNT-induced FZD-DVL dynamics, we made use of three DEP mutants previously reported to interfere with FZD-DVL recruitment and downstream WNT/β-catenin signaling^3,6-9^. The DEP G436P and K446M mutants were both recruited to FZD_5_ albeit with a slightly reduced affinity (represented by BRET_50_) for FZD_5_ (Fig. 5E). Thus, the mutations affect FZD-DEP binding suggesting that the finger loop (K446M) itself and the region at its base (G436P) are important for the FZD-DEP interface. The DEP L445E mutant, which is part of the finger loop in the DEP domain, was not recruited to FZD_5_, further corroborating the importance of the DEP finger loop for FZD_5_-DEP interaction. Interestingly, microscopy-based analysis indicated that the DEP K446M mutant does not bind to FZD_5_ (*11*); the more quantitative approach of BRET analysis, however, was able to detect the weakened interaction of this DEP mutant with its receptors, clearly underlining the advantageous dynamic range of the BRET-based approach (Fig. 5C).

With regard to the WNT-induced kinetics, we observed a drastic reduction of the ΔBRET_max_ response observed between FZD_5_ and DEP K446M compared to wildtype DEP in response to both WNT-3A and WNT-5A. Most importantly, the efficacy differences of WNT-3A and WNT-5A observed for the FZD-DEP conformational dynamics were abrogated in the experiments performed with mutant DEP K446M (Fig. 6D). Thus, our data suggest that the finger loop is not only important for basal FZD-DEP interaction but also central for WNT-induced FZD_5_-DEP dynamics. Regarding the G436P DEP mutant, we observed a higher efficacy for WNT-3A compared to WNT-5A (Fig. 6D). Thus, this DEP mutation affects the FZD-DEP interaction in a WNT-selective manner arguing that WNT-3A and WNT-5A stabilize distinct FZD conformations that show more or less efficient DEP interaction, respectively. Furthermore, the DEP G436P mutation increased the rate constants (Fig. 6C) for the WNT-induced BRET changes, suggesting that this structural change in the DEP domain evoked by the mutation is either involved in FZD interaction or that DEP dimerization could present a rate-limiting step affecting agonist-induced conformational kinetics (*11*). In summary, our mutational analysis allowed us to distinguish between constitutive FZD-DEP interaction, WNT-induced conformational dynamics and WNT-selective processes as part of an agonist-induced functional selectivity in a WNT-FZD_5_-DEP complex.

Based on the current working model of WNT/β-catenin signaling, WNTs bind FZD and LRP5/6 to allow for recruitment of DVL and formation of the signalosome complex (*30*). We tested this model and concluded that WNTs induce FZD-DVL conformational dynamics independently of LRP5/6, clearly suggesting that (i) WNT-FZD interactions can occur independently of LRP5/6, that (ii) agonist-induced FZD-DVL dynamics potentially precede LRP5/6 signalosome formation and that (iii) WNT-induced conformational changes of FZDs regulate the initial communication with DVL in a receptor complex reminiscent of the ternary complex described for agonist- and effector-bound GPCRs (*4, 22*). The emerging concept of conformational dynamics of FZDs is in agreement with general concepts of GPCR activation (*41*) and is further supported by previously reported WNT concentration response curves for fluorescence changes in cpGFP-tagged FZD conformational sensors (*3*) and the WNT-induced dynamics of the cysteine-rich domain of FZDs that precede alterations in core conformation (*42*). Thus, WNT-induced pathway selection and signal initiation of FZDs is not only determined by their proximity to different co-receptors but appears to be – at least in part – determined by a conformational selection of WNT-FZD-DVL interaction.

In summary, the data presented here support the notion that WNT binding to FZD_5_ results in conformational dynamics of the FZD-DVL interaction supportive of a cooperative, allosteric interaction in a WNT-FZD-DVL complex. These conformational FZD_5_-DVL dynamics are DVL oligomerization-independent, and the evidence points to a ligand-selective FZD conformation-driven process at the interface between FZD and the DEP finger loop. While this concept is supported by the current and previously published data, additional structural insight is required to understand how conformational rearrangement of the WNT-FZD-DVL complex initiates downstream signaling and how its function is integrated with ligand-induced signalosome formation and the formation of WNT-FZD-G protein complexes (*4*).

## MATERIALS AND METHODS

### Cell culture

For all experiments, ΔFZD_1-10_ HEK293T cells were used if nothing else was stated. ΔFZD_1-10_ HEK293T cells (*43*) and ΔLRP5/6 HEK293 T-Rex cells (*44*) were cultured in DMEM (HyClone SH30081) supplemented with 10% fetal bovine serum and 1% penicillin/streptomycin (Thermo Fisher Scientific 10270106 and 15140) in a humidified incubator at 37 °C and 5% CO_2_. All cell culture plastics were from Sarstedt unless stated otherwise. For cell transfection, ΔFZD_1-10_ or ΔLRP5/6 cells were transiently transfected in suspension using Lipofectamine 2000 (Thermo Fisher Scientific 11668). An approximately 80-90% confluent T-75 flask was re-suspended in 20 ml yielding a suspension with 4-5×10^5^ of cells ml^-1^. A total of 1 µg of DNA was used to make 1 ml of transfected cell suspension and was always filled up with empty pcDNA3.1 plasmid. Plasmid amounts used in a transfection are later presented as percentage (%) of total plasmid DNA used in that transfection, e.g. 2% of Nluc plasmid DNA is defined as 20 ng of Nluc plasmid DNA used to transfect 4-5×10^5^ of cells in 1 ml.

### Cloning and plasmids

Venus-DVL2 was subcloned from FLAG-DVL2 (Addgene #24802) into pVenus-C1 with HindIII and BamHI. HA-FZD_5_-Nluc was cloned using prolonged-overlap-extension (POE) PCR techniques. ΔDEP Venus-DVL2 was cloned with the Q5 Site-Directed Mutagenesis kit (NEB #E0554S) using Venus-DVL2 as a template. Venus-DVL2-M2/M4 was generated using the GeneArt Site-directed mutagenesis kit (Thermo Fisher Scientific). DEP-Venus was cloned from a gBlock (Supplementary Information text) into mVenus-N1 (Addgene #27793) using HindIII and BamHI restriction sites. The β_2_AR-Nluc construct was cloned by fusing Nluc to the C-terminus of FLAG-SNAP-β_2_AR using XbaI and NotI restriction sites and DNA fragment ligation. Nluc-DVL2 was generated and validated previously (REF to Mol switch paper). HA-FZD_5_ was from Thomas Sakmar and Venus-KRas was from Nevin Lambert.

### Ligands

Recombinant human WNT-3A and human/mouse WNT-5A were from RnD Systems/Biotechne (#5036-WN, #645-WN). WNTs were dissolved to 100 µg/ml in filter-sterilized 0.1% BSA in PBS (HyClone) and stored at 4 °C. WNTs were diluted in filter-sterilized 0.1% BSA in HBSS (HyClone) and vehicle control was diluted in filter sterilized 0.1% BSA in HBSS solution and used for the serial dilution. WNTs were kept on ice when handled. DKK1 was mixed with the WNT solution before stimulation. For experiments with inactivated WNT-3A, the WNT-3A dilution and the corresponding vehicle control were subjected to a heat-freeze cycle (1 h at 65 °C and 1 h at −20 °C) twice before usage in the experiments. Digitonin was purchased from Sigma (#D141), dissolved at 10 mg/ml in DMSO and stored at −20 °C.

### BRET

For all BRET experiments, 2% of Nluc plasmid was used if nothing else was stated. In the BRET titration experiments, up to 75% of Venus plasmid was used and for Venus-DVL2 and Venus-DVL2-M2/M4 a minimum of 1.7% and for DEP-Venus 0.1% plasmid was used, as well as one condition without Venus plasmid resulting in a total of 8 conditions. In the stimulation experiments, 50% of Venus plasmid was used. 100 µl of cells were seeded into poly-D-lysine (PDL)-coated white 96-well plates with flat bottom (Greiner BioOne). 22-26h post transfection, cells were washed once with 200 µl of HBSS and then kept in HBSS. In the titration experiments, Venus fluorescence was measured first and subsequently 10 µl of Coelenterazine h (Biosynth C-7004) (2.5 µM final concentration) were added yielding a final volume of 100 µl. 5 min after addition, luminescence was read 3 times. In the stimulation experiments, Venus fluorescence was first measured and subsequently 10 µl of Furimazine (Promega) (final dilution of 1:1000) were added to a final volume of 90 µl and 5 min after addition luminescence was read 3 times. Subsequently, 10 µl of ligand were added to a final volume of 100 µl and luminescence reading was continued 28 times for a total of 62 min. All measurements were performed at 37 °C. Fluorescence was measured using a ClarioStar (BMG) plate reader (497-15 excitation and 540-20 emission). The BRET ratio was determined as the ratio of light emitted by Venus (acceptor) and light emitted by Nluc (donor). Net BRET was calculated by subtracting the donor only (transfection without Venus) BRET ratio. ΔBRET was calculated as the difference between vehicle and WNT-treated wells, where each well was normalized to the mean value of the first three reads. The BRET emission signal was measured using a ClarioStar (BMG) plate reader with bandpass filters 535-30 nm (acceptor) and 475-30 nm (donor). For experiments with inactivated WNT-3A, cells were transfected with 1% of FZD_5_-Nluc plasmid and 50% of Venus plasmid (DEP-Venus or Venus-DVL2). Experiments were run two days after transfection following the protocol described above using a TECAN Spark multimode reader. Emission intensity of the donor (Nluc) was detected with a 445-470 nm bandpass filter and the acceptor emission intensity (Venus) with a 520-545 nm bandpass filter. An integration time of 0.1 s was applied for recording of both emissions. In the bystander BRET experiments, 78% of Venus-KRas was used together with 20% of HA-FZD_5_. 24h post-transfection, the cells were washed with HBSS and kept in this buffer with 1:1000 furimazine. Basal BRET (460-500 nm, 520-560 nm; 0.2 s integration time) was measured three times on TECAN Spark. Subsequently, vehicle or WNT-3A or WNT-5A (at a final concentration of 1000 ng/ml) were added and BRET was measured for 60 min (47 cycles). In the experiments with digitonin, this detergent (10 μg/ml final concentration) or vehicle were added and BRET was sampled 24 times during 30 min using TECAN Spark.

### ELISA surface expression

For quantification of cell surface expression of the different receptor constructs, cells were transfected with 50% of the indicated receptor plasmid. 100 µl of cell suspension were seeded onto a transparent PDL-coated 96-well cell culture plate with solid flat bottom. 24 h later the medium was dispensed from the wells. Cells were washed once with 200 µl of ice-cold wash buffer (0.5% BSA in PBS), and incubated on ice with 25 µl of primary antibody solution (1% BSA in PBS with either anti-HA 1:500 (Abcam ab9110) or anti-FLAG 1:500 (Sigma-Aldrich F1804)) for 1h. Subsequently, cells were washed as described above four times and then incubated on ice with 50 µl of secondary antibody solution (1% BSA in PBS with either HRP-conjugated anti-mouse 1:5000 (Invitrogen 31430) or anti-rabbit 1:5000 (Invitrogen 31460)) for 1 h, after which cells were washed four times. Lastly, 50 µl of TMB solution (3,3′,5,5′-Tetramethylbenzidine, Sigma T0440) were added to each well and incubated for 20 min after which 50 µl of 2M HCl were added and incubated for an additional 20 min. Absorbance (450 nm) was read with a POLARstar Omega plate reader (BMG).

### TCF/LEF luciferase reporter assay (TOPFlash)

ΔFZD_1-10_ cells were transfected in suspension with 2% FZD_5_-Nluc, 25% M50 Super 8xTOPFlash (Addgene #12456) and 5% pRL-TK Luc (Promega E2241) plasmid and 100 µL of the transfected cell suspension were seeded into a white PDL-coated 96-well flat bottom plate (Nunc). 24 h after transfection, cells were washed once with 200 µl dPBS and medium was changed to DMEM without FBS containing either vehicle control, 300 ng/ml of WNT-3A or 300 ng/ml of WNT-3A together with 300 ng/ml of DKK1. 24 h post stimulation, cells were washed once with 200 µl of dPBS and subsequently analyzed using a Dual-Luciferase Reporter Assay System kit (Promega E1910) according to the manufacturer’s instructions with the following modifications: 20 µl of passive lysis buffer, 20 µl of LARII and 20 µl of Stop & Glo reagent were used. The luminescence was measured using a CLARIOstar microplate reader (BMG) (Firefly and Renilla were measured at 580 ± 40 nm and 480 ± 40 nm, respectively).

### Data analysis and statistics

All data were analyzed with GraphPad Prism 8 (San Diego, CA, US). Data points for the titration experiments represent mean ± SD. Data points for the kinetic experiments, concentration response curves and TOPFlash represent mean ± SEM. Concentration response curves were plotted as the response 30 min after stimulation from each concentration of the kinetic experiments. The concentration response curves were fit using a non-linear three parameters model selected based upon an extra sum-of-squares F test in comparison to a four parameters model (P < 0.05). For non-saturating concentration response curves, caution should be taken when interpretating the EC_50_ values, which are provided, when the software (GraphPad Prism 8) allowed their definition. The TOPFlash statistical test was done using one-way ANOVA with Dunnett’s post hoc analysis. The statistical differences for the surface expression and basal recruitment of DVL2 in the bystander BRET assays were tested with paired Student’s t-test (** P < 0.01, *** P < 0.001). WNT-induced kinetics were analyzed with a plateau followed by one phase association equation, data points represent mean ± SD. Individual rate constants from each experiment were compared with a two-way ANOVA with Fisher’s least significant difference post hoc analysis.

## Acknowledgments

Thank you to Anna Krook for access to the CLARIOstar plate reader, to Benoit Vanhollenbeke for the ΔFZD_1-10_ HEK293T cells, to Nevin A. Lambert for assistance with gBlock design, Vitezslav Bryja for the ΔLRP5/6 HEK293 T-rex cells and Mariann Bienz for the ΔDVL1-3 HEK293 T cells. Figure illustrations were created with BioRender.com.

## Author contributions

Conceptualization: CFB, GS Methodology: CFB

Investigation: CFB, PK, LG, MKJ, HS

Visualization: CFB

Funding acquisition: CFB, PK, GS Project administration: CFB, GS Supervision: GS

Writing – original draft: CFB, GS

Writing – review & editing: CFB, GS, PK, LG

## Data and materials availability

All data are available in the main text or the supplementary materials.

## Supplementary Information

### Supplementary Materials and Methods

#### DEP gBlock

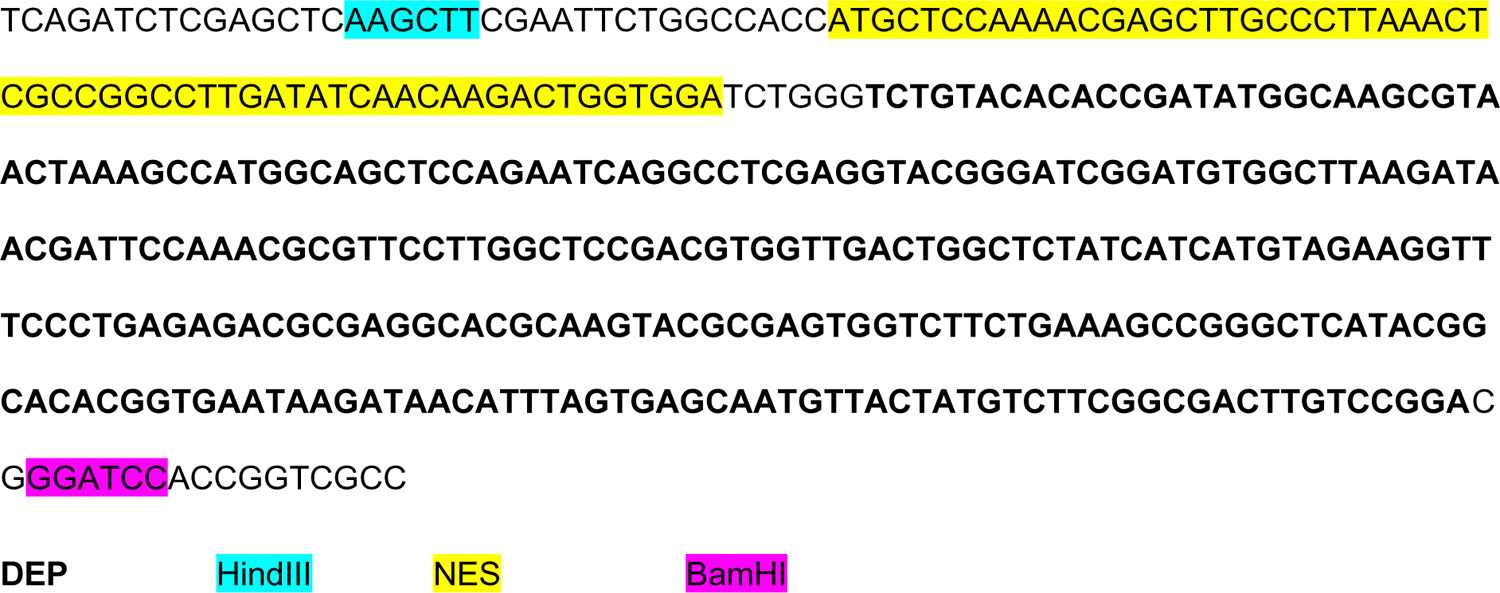

#### HA-FZD_5_-Nluc prolonged overlapping extension primers

Vector forward: 5’ GAACGCATTCTGGCGTAAGTACCGCCTCCTCGGATG

Vector reverse: 5’ ATCTTCGAGTGTGAAGACGACGTGGCTCAGAGACA

Insert forward: 5’ TGTCTCTGAGCCACGTCGTCTTCACACTCGAAGATTTCG

Insert reverse: 5’ CATCCGAGGAGGCGGTACTTACGCCAGAATGCGTTC

#### ΔDEP Venus-DVL2

Vector forward: 5’ GGCTGTGAGAGTTACCTAGTTAACCTC

Vector reverse: 5’ GAGACCCCGGCCTTCGCA

**Fig. S1.**
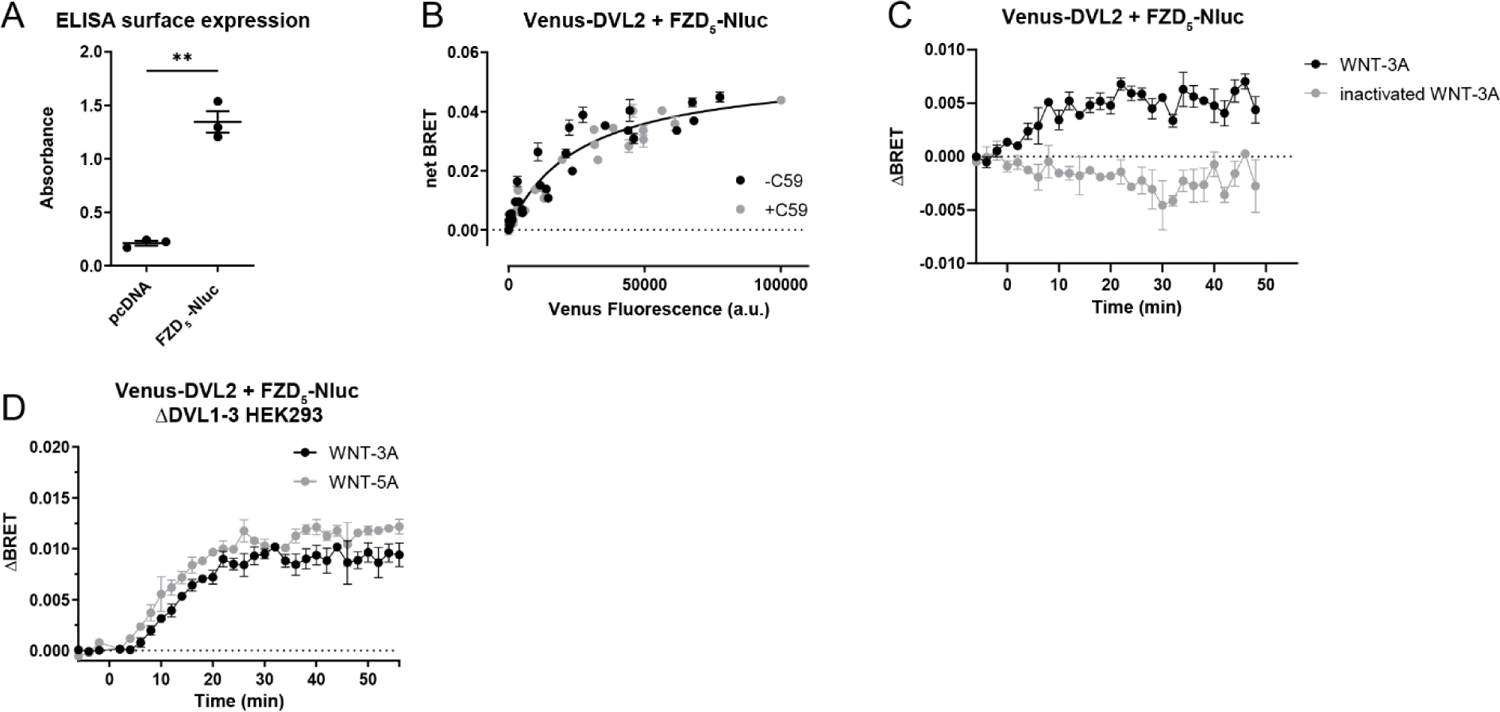
Validation of the FZD_5_-DVL2 BRET approach. (**A**) ELISA surface expression of HA-FZD_5_-Nluc in comparison to pcDNA. Data are presented as mean ± SEM of 3 independent experiments and were analyzed using ratio paired Student’s t-test (** P < 0.01). (**B**) Venus-DVL2 was titrated with a fixed amount of Nluc-tagged FZD_5_ treated with either vehicle or 10 nM C59 overnight to investigate the dependence of basal recruitment of DVL to FZDs on the secretion of endogenously expressed WNTs. The curve fit was calculated using One-site specific binding comparing fits using extra sum-of-squares F test (P < 0.05), and data are presented as mean ± SD for 4 independent experiments. (**C**) The kinetic BRET response was monitored between Venus-DVL2 and FZD_5_-Nluc (Venus:Nluc ratio 50:1) in ΔFZD_1-10_ cells with stimulation of either WNT-3A (1000 ng/ml) or inactivated WNT-3A (1000 ng/ml). Data are presented as mean ± SEM of 3 independent experiments. (**D**) Fixed amounts of Venus-DVL2 and Nluc-tagged FZD_5_ (Venus:Nluc ratio 25:1) were transfected into ΔDVL1-3 cells to investigate the potential role of endogenous DVL for the observed FZD_5_-DVL dynamics upon WNT stimulation. The kinetic BRET response between Venus-DVL2 and FZD_5_-Nluc was monitored with stimulation of either 1000 ng/ml WNT-3A or WNT-5A. Data are presented as mean ± SEM of 3 independent experiments.

**Fig. S2.**
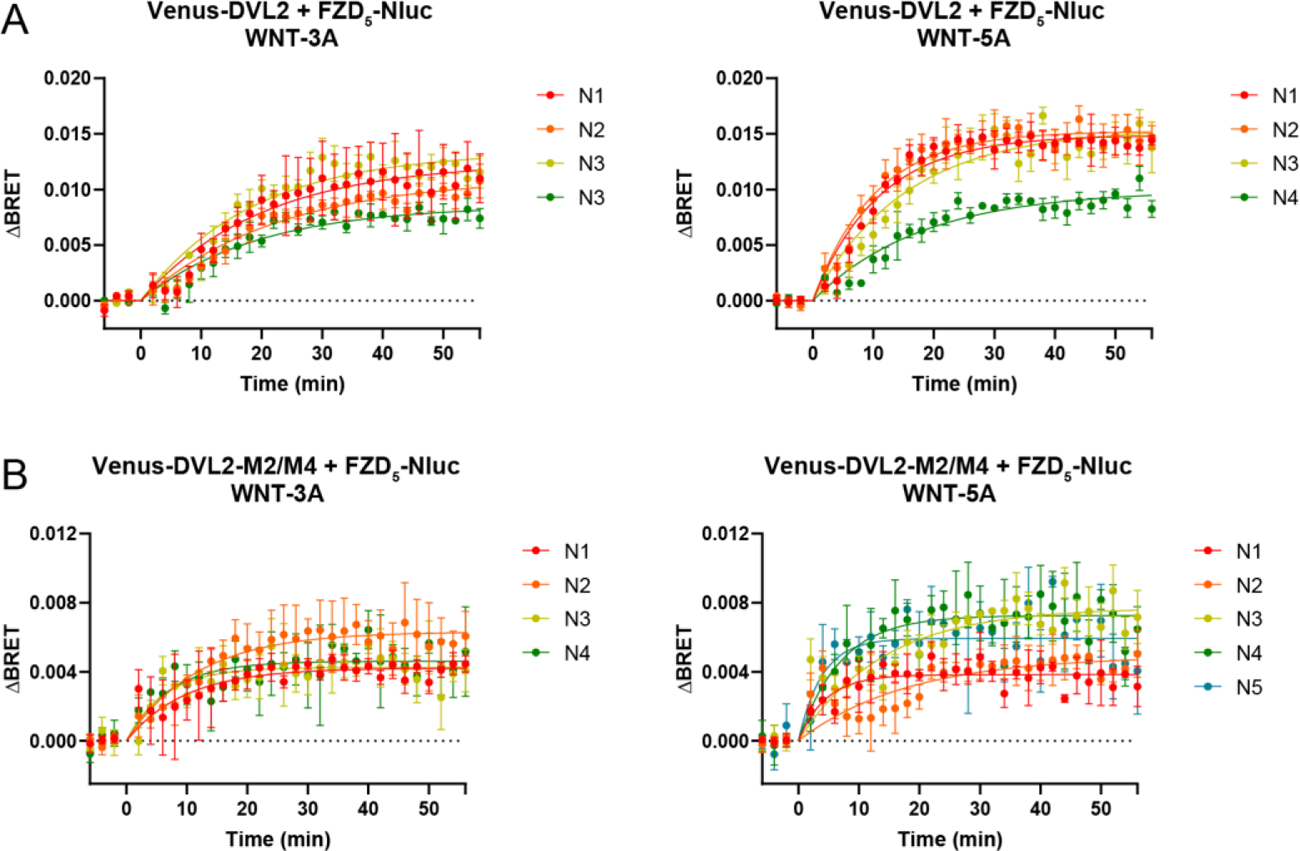
Kinetic analysis of the WNT-induced conformational dynamics of FZD_5_-DVL2 interaction. (**A-B**) The kinetic BRET responses were monitored between (A) Venus-DVL2 or (B) DVL2-M2/M4 mutant and FZD_5_-Nluc (Venus:Nluc ratio 25:1) in ΔFZD_1-10_ cells with stimulation of either WNT-3A (1 µg/ml) or WNT-5A (1 µg/ml). Graphs show four to five independent experiments (N1-N5) with means ± SD. Data were fitted using the plateau followed by one phase association equation.

**Fig. S3.**
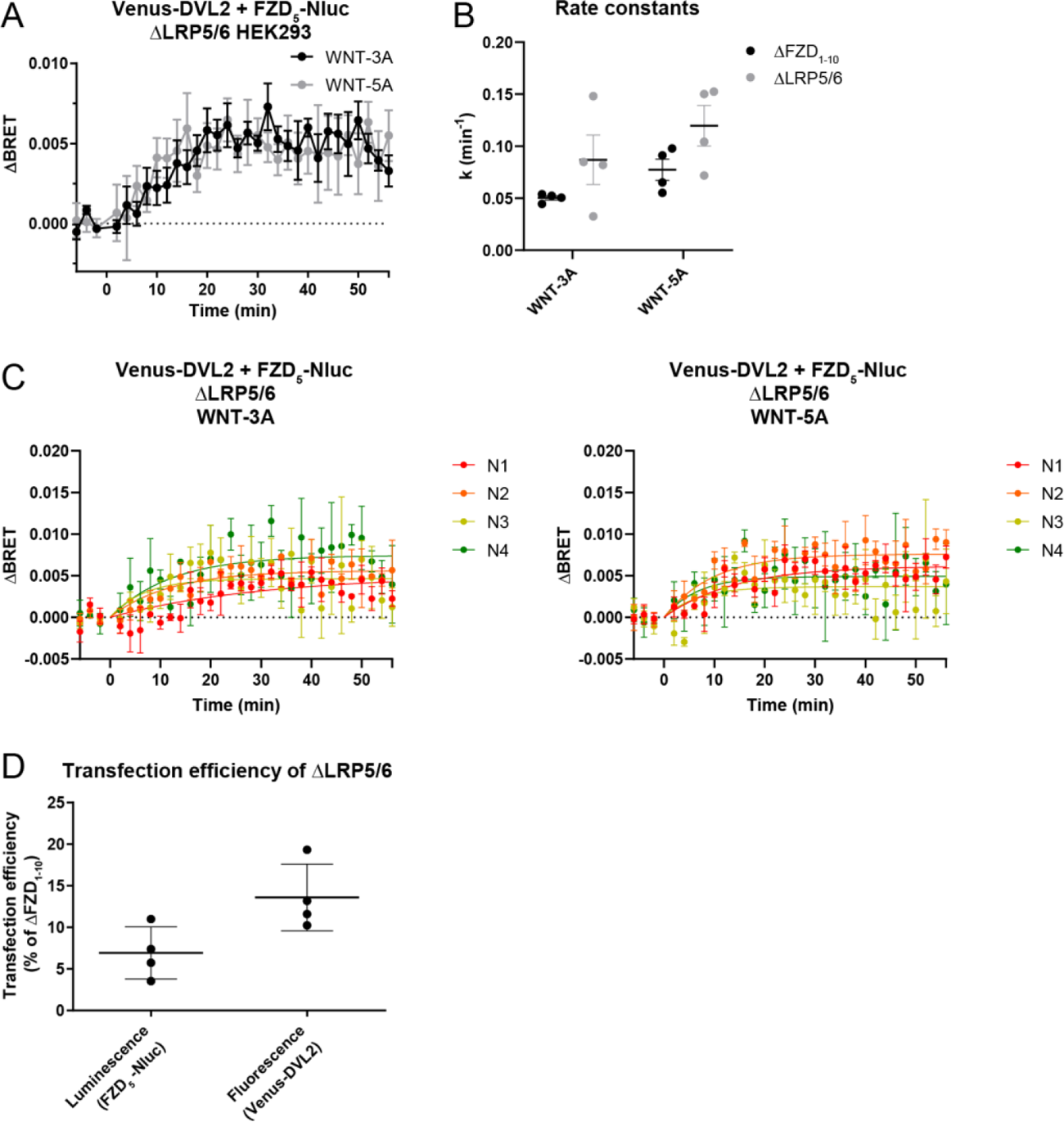
WNT-induced FZD_5_-DVL2 conformational dynamics is independent from LRP5/6. (**A**) The kinetic BRET response was monitored between Venus-DVL2 and FZD_5_-Nluc (Venus:Nluc ratio 25:1) in ΔLRP5/6 cells during stimulation with 1 µg/ml of WNT-3A or WNT-5A. Data are presented as mean ± SEM of four independent experiments. (**B**) Comparison of Venus-DVL2 rate constants, k, in ΔFZD_1-10_ and ΔLRP5/6 cells upon WNT stimulation. Data were extracted from Fig. S2A and S3C. No statistically significant difference was found between ΔFZD_1-10_ and ΔLRP5/6 cells (two-way ANOVA with Fisher’s LSD post hoc analysis). (**C**) Kinetic fits for WNT-induced conformational dynamics of DVL2-FZD_5_ in ΔLRP5/6 cells. Graphs show four independent experiments (N1-N4) with means ± SD. Data were fitted using the plateau followed by one phase association equation. (**D**) The transfection efficiency of ΔLRP5/6 cells was compared to that of ΔFZD_1-10_ cells. Luminescence and fluorescence values are from the wells stimulated with 1 µg/ml of WNT in experiments from Fig. 1C (ΔFZD_1-10_) or from wells stimulated with WNTs in Fig. S3A (ΔLRP5/6). Data are presented as mean ± SEM of four independent experiments.

**Fig. S4.**
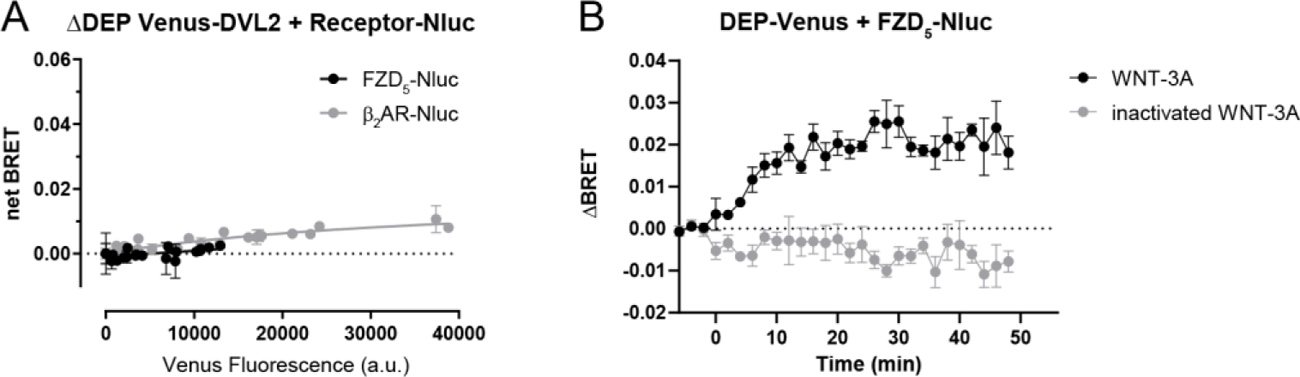
Validation of the direct FZD_5_-DEP BRET approach. (**A**) The DEP domain of DVL2 is essential for basal FZD_5_-DVL recruitment. ΔDEP Venus-DVL2 was titrated with a fixed amount of Nluc-FZD_5_ in ΔFZD_1-10_ cells to assess basal recruitment of ΔDEP DVL2 to FZD_5_; β_2_AR was used as negative control. net BRET is presented as mean ± SD of three independent experiments. (**B**) The kinetic BRET response was monitored between DEP-Venus and FZD_5_-Nluc (Venus:Nluc ratio 50:1) in ΔFZD_1-10_ cells with stimulation of either WNT-3A (1 µg/ml) or inactivated WNT-3A (1 µg/ml). Data are presented as mean ± SEM of three independent experiments.

**Fig. S5.**
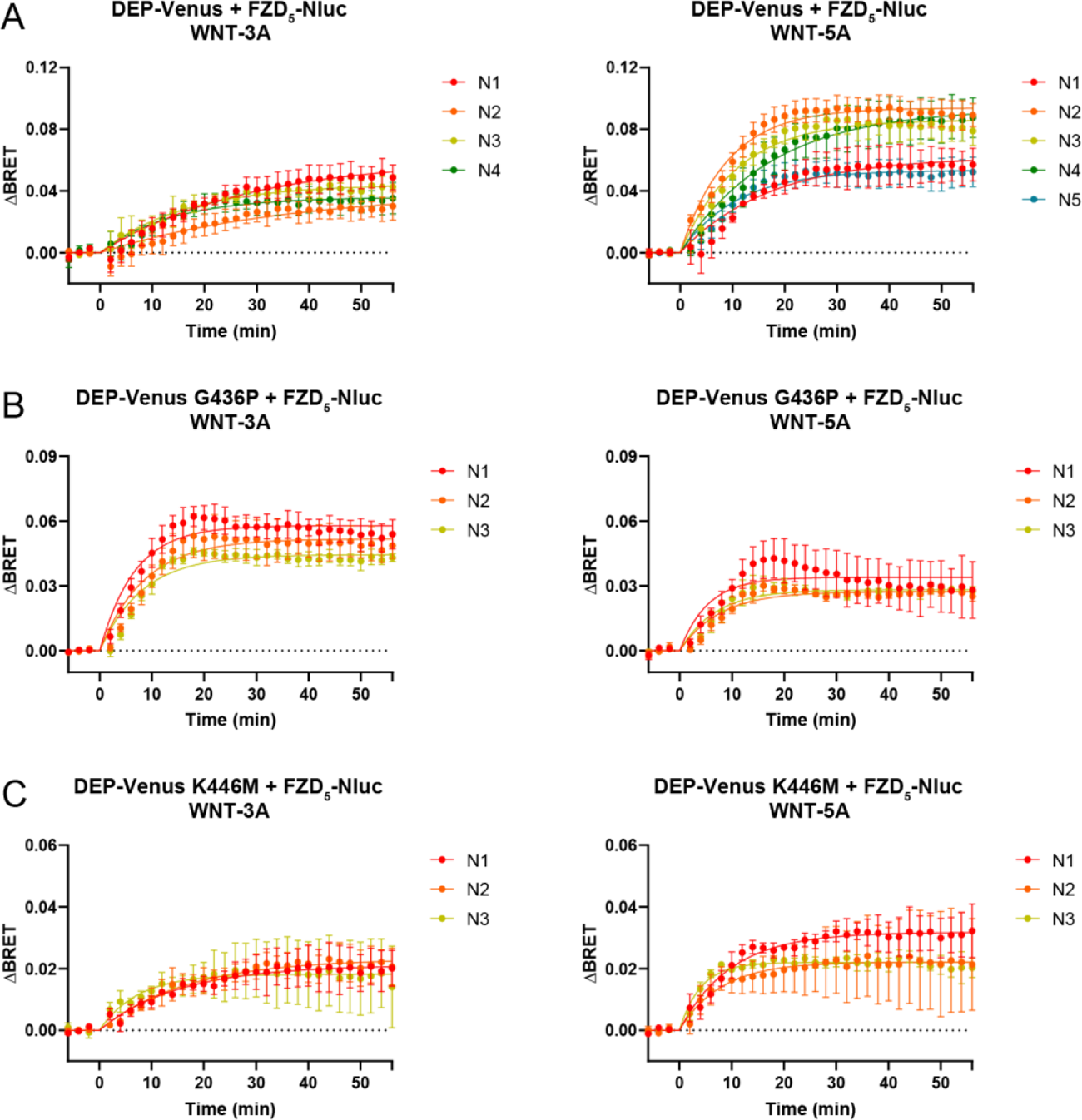
Kinetic analysis of WNT-induced conformational dynamics of FZD_5_-DEP interaction. (**A-C**) The kinetic BRET responses were monitored between DEP-Venus, its respective mutants and FZD_5_-Nluc (Venus:Nluc ratio 25:1) in ΔFZD_1-10_ cells with stimulation of either WNT-3A (1 µg/ml) or WNT-5A (1 µg/ml). Graphs show three to five independent experiments (N1-N5) with means ± SD. Data were fitted using the plateau followed by one phase association equation.

